# Severe acute dehydration in a desert rodent elicits a transcriptional response that effectively prevents kidney injury

**DOI:** 10.1101/103077

**Authors:** MacManes Matthew

## Abstract

Animal living in desert environments are forced to survive despite severe heat, intense solar radiation, and both acute and chronic dehydration. Indeed, these animals have evolved phenotypes that effectively address these environmental stressors. To begin to understand the ways in which the desert adapted rodent *P. eremicus* survives, we performed an experiment by which we subjected reproductively mature adults to profound acute dehydration, during which they lost on average 23% of their body weight. Animals react via a series of changes in the kidney, which include modulating expression of genes responsible for reducing the rate of transcription, and maintaining water and salt balance. Extracellular matrix turnover appears to be decreased, and apoptosis is limited. Serum Creatinine and other biomarkers of kidney injury are not elevated, which is different than the canonical human response, suggesting that transcriptional changes caused by acute dehydration effectively prohibit widespread kidney damage in the cactus mouse.

## Introduction

Dehydration, whether caused by exposure to extreme environmental conditions, water deprivation, or by infection (*e.g.*, diarrheal illnesses), represents a significant threat to human life (1). In spite of modern medicine, millions of people die every year from dehydration (2). Compounding issues of exposure and illness are public health issues regarding the delivery of safe drinking water. With global climate change, it is expected that these challenges will only become more severe (3). As a result, research providing insight into the ecological genomics underlying renal homeostasis and physiologic resistance to acute and chronic dehydration is urgently needed. The response to severe acute dehydration in humans and traditional mammalian models is generally limited and may fail to prevent severe electrolyte imbalance, renal impairment or failure, or even death (4–7). In contrast, desert-living mammals have evolved phenotypes making them adept at surviving acute and chronic dehydration (8, 9). These phenotypes include numerous aspects of behavior (*e.g.*, many animals are nocturnal or crepuscular), morphology (*e.g.*, a nasal countercurrent system (10)), physiology (*e.g.*, renal histology (11–14), and metabolic water production (9, 15).

In addition to the mechanisms described above, desert animals are thought to possess specialized machinery allowing for the efficient maintenance of water and solute balance, in spite of the obvious challenges presented to them as part of desert living. The aquaporins, a large protein family integral to the maintenance of water balance in the kidney (16–18) were the first genes implicated in desert rodents’ remarkable abilities (19), and immediately became canonized as one of the critical adaptation required by at least some desert rodents. To this end, several other studies provided evidence of the import of this gene family (20–23), which in brief, allows water to be resorbed from the urinary lumen back into circulation, thereby reducing water loss. With regards to solute balance, several studies have found members of another gene family - the Solute Carriers - to be differentially expressed or under lineage-specific positive selection (20, 24). Broadly speaking, these transport molecules are responsible for the maintenance of electrolyte levels requisite for the normal function of virtually all bodily processes (25).

Dehydration and the resultant hyperosmolar state, the direct result of intense heat and aridity, is typically coupled with electrolyte derangement (26, 27) and therefore maintaining electrolyte gradients requisite for proper function may be particularly challenging for desert animals. Given these physiological challenges, specialized function of the Aquaporins and Solute Carriers may provide at least partial mechanisms thereby allowing for rodents to survive in desert-conditions.

The cell-level consequences of failure of these mechanisms (*e.g.*, salt and water imbalance) is well characterized, at least for non-desert adapted animals (27–29). In brief, dehydration results in a hyperosmotic state, which, amongst other things, results in cell cycle arrest (30), double strand DNA breaks (31) and disruption of repair mechanisms (32), oxidative stress (33), inhibition of transcription and translation (34). Additionally, hyperosmolarity is known to inhibit protein folding in the Endoplasmic Reticulum (ER - 1, 35). Inhibited protein folding results in physical crowding of the ER, and invocation of the unfolded protein response (UPR), which provides mechanisms for either restoring homeostasis, or in cases where stress is too severe or prolonged, facilitating apoptosis (36). Given this, the response to a hyperosmolar environment is profound, with disruptions to vital cellular processes. Here, we hypothesize that these responses will be less profound in an animal adapted to exceptionally dry conditions. In particular, we predict that while cellular stress will be apparent, signs of widespread apoptosis and tissue damage will not. To this end this study seeks to provide transcriptomic evidence to test this hypothesis.

To better understand the effects of dehydration in desert-adapted animals, and in particular the cellular effects, we conducted an experiment whereby we exposed a set of animals (*Peromyscus eremicus*) to experimental dehydration. The rodent *P. eremicus* (cactus mouse) represents an ideal model in which to study the genomic architecture of adaptive dehydration-resistance, and that of acute dehydration. These animals have evolved in the extremely hot and dry desert regions of the US Southwest (37). They survive their entire lives without drinking water, and in many cases urinating. Though highly adapted to desert environments, *P. eremicus* retains many of the useful characteristics of more well established rodent models, including the facility with which they reproduce in the lab (38) and the availability of multiple sequenced genomes (*P. maniculatus, P polionotus, P. californicus*). In addition, the genus *Peromyscus* - known as the *Drosophila* of North American Mammalogy (39), has been the focus of ecological study for decades (24, 38, 40–43) and has recently emerged as a powerful clade of organisms within which to study the genetics of adaptive phenotypes (43–45).

## METHODS

### Animal Care and Experimental Model

The cactus mice used for this study include only captive born individuals purchased from the *Peromyscus* Genetic Stock Center (Columbia, South Carolina). The animals, originally collected from a hot-desert location in Arizona, have been housed for several generations at the University of New Hampshire in conditions that mimic temperature and humidity levels in southwestern US deserts, as described previously (46). Animals were randomly assigned to either a dehydration treatment group (n=19) or a control group (n=19). Mice that were assigned to the dehydration group were weighed and then water deprived for ~72 hours directly prior to euthanasia. All mice were again weighed prior to sacrifice. Individuals in the study were collected at approximately noon (6 hours after lights on), and when exposed to 90°F for four hours, between September 2014 – April 2016.

Cactus mice were sacrificed via isoflurane overdose and decapitation in accordance with University of New Hampshire Animal Care and Use Committee guidelines (protocol number 130902) and guidelines established by the American Society of Mammalogists (47). Trunk blood samples were collected following decapitation for serum electrolyte analyses with an Abaxis Vetscan VS2 using critical care cartridges (Abaxis). The complete methodology and results of the electrolyte study, as well as the reported measures of water consumption and weight loss due to dehydration are described fully elsewhere (Kordonowy et al., 2016). Both kidneys were harvested within five minutes of euthanasia, placed in RNAlater (Ambion Life Technologies), flash-frozen in liquid nitrogen, and stored at −80° degree Celsius. A TRIzol, chloroform protocol was implemented for RNA extraction (Ambion Life Technologies). The quantity and quality of the RNA product was evaluated with a Qubit 2.0 Fluorometer (Invitrogen) and a Tapestation 2200 (Agilent Technologies, Palo Alto, USA).

### Illumina Library Preparation and Sequencing

Tissues frozen in RNALater were thawed on ice in an RNAse-free work environment. Total RNA was extracted using a standard Trizol extraction protocol (Thermo Fisher Scientific, Waltham, MA). The quality of the resultant extracted total RNA was characterized using the Tapestation 2200 Instrument (Agilent, Santa Clara, CA), after which Illumina sequence libraries were prepared using the TruSeq RNA Stranded LT Kit (Illumina). The Tapestation 2200 Instrument was, again, used to determine the quality and concentration of these libraries. Each library was diluted to 2nM with sterile ddH_2_O, and pooled in a multiplexed library sample. The multiplexed library sample was then sent to the New York Genome Center for 125 base pair paired-end sequencing on a HiSeq 2500 platform.

### Sequence Quality Control and Assembly

Sequence read data were downloaded and quality checked using FastQC version 0.11.5 (48). A *de novo* transcriptome for the cactus mouse kidney was assembled following the Oyster River Protocol for Transcriptome Assembly (49). In brief, 59 million paired end reads from two individuals (1 control and 1 dehydrated) were error corrected using RCorrector version 1.0.2 (50). Adapters, as well as bases with a Phred score <2 were trimmed using Skewer 0.2.2 (51), and assembly was carried out using Trinity version 2.2.0 (52), Binpacker version 1.0 (53), and Shannon version 0.0.2 (54), after which the resultant transcriptomes were merged using the software package transfuse version 0.5.0 (https://github.com/cboursnell/transfuse). Lowly expressed transcripts, defined as those with an abundance of less than one transcript-per-million (TPM<1), were filtered out of the dataset. The resultant assembly was annotated using the software package dammit (https://github.com/camillescott/dammit), and evaluated using BUSCO 2.0 (55) and TransRate version 1.0.1 (56).

### Mapping and Global Analysis of Differential Gene Expression

After quality and adapter trimming to a Phred score =2, reads were quasimapped to the cactus mouse kidney transcriptome after an index was prepared using Salmon 0.7.2 (57). Kidney transcript IDs were mapped to genes from *Mus musculus* proteome version GRCm38.p5, using BLAST (58). All data were then imported into the R statistical package (version 3.3.0) (59) using tximport (60) for gene level evaluation of gene expression, which was calculated using edgeR (version 3.1.4) (61) following TMM normalization and correction for multiple hypothesis tests by setting the false discovery rate (FDR) to 1%. Gene ontology enrichment analysis was carried out using the Kolmogorov-Smirnov test (62) for significance in the R package, topGO (63).

### Coexpression network analysis

TMM normalized count data for each individual, as well as physiology data consisting of individual water intake, weight loss and electrolyte data (described in [(64)], and available as Supplementary dataset 1, name = phys4.csv on GitHub) were imported into the package WCGNA (65). Gene co-expression modules were described, then a heatmap that describes the relationship between co-expression modules and phenotype was produced. Gene ontology terms for genes residing in specific modules were generated.

## Results and Discussion

### Animal Response to Acute Dehydration

Thought the response to dehydration is fully described elsewhere (64), in brief, animals lost on average 23.2% of their body weight (median=23.9%, SD=5.3%, min=12.3%, max=32.3%). Accordingly, serum electrolytes (Sodium, Calcium, Bicarb) were elevated in dehydrated animals, as well BUN. Interestingly, a serum marker of kidney damage, Creatinine, is not.

### Sequence Read Data & Code Availability

In total, 38 kidneys (19 dehydrated, 19 control) were sequenced. Each sample was sequenced with between 2.3 million and 60.6 million read pairs (mean=16.8M, SD=1.3M). All read data are available using the project ID PRJEBXXX. Code used for the analysis of these data are available at https://git.io/vMN6l.

### Transcriptome Assembly and Evaluation

The kidney transcriptome assembly consists of 79,660 transcripts. This assembly, along with the annotations contained in gff3 format, is available at (dryad on acceptance). The transcriptome contains 24,794 transcripts with at least 1 hit to the Pfam database, 59,733 hits to OrthoDB (66), 71,857 hits to Uniref90, 2,831 hits to the transporter database (67), and 924 hits to Rfam (68). These 79,660 assembled transcripts map to 13,697 unique genes in the *Mus musculus* genome. The evaluation using BUSCO and TransRate are presented in Table 1 and 2 respectively.

**Table 1:** Quality control statistics for the assembled transcriptome: BUSCO metrics include statistics regarding the number of universal single copy orthologs found in the assembly. 78% of the Mammalian BUSCOs were identified as full length (complete) sequences, while 5.4% were found to be fragmented. 30.3% were found in greater than 1 copy in the cactus mouse kidney transcriptome.

|  | BUSCO |  |  |
| --- | --- | --- | --- |
|  | Complete | Duplicated | Fragmented |
| PEER Kidney 1.0.0 | 78% | 30.3% | 5.4% |

**Table 2:**
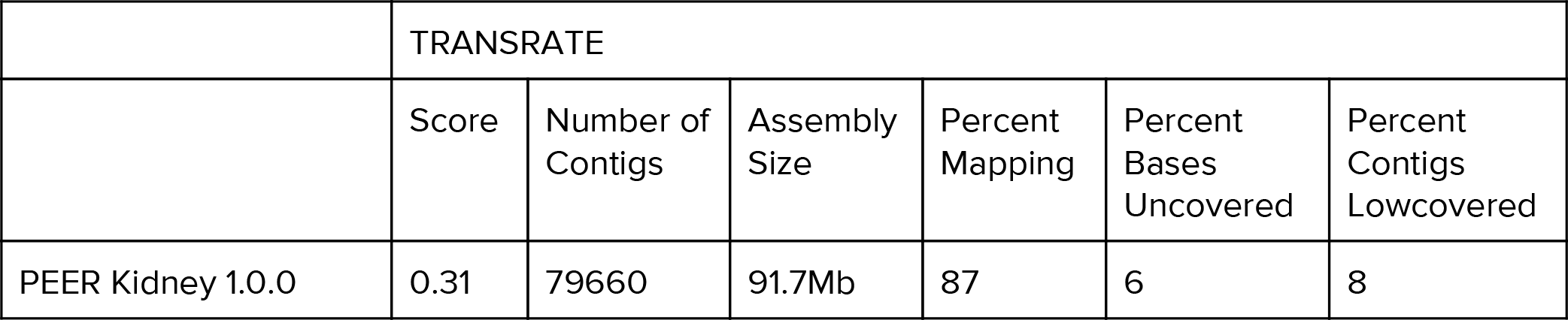
Quality control statistics for the assembled transcriptome: TransRate metrics are derived from mapping RNAseq reads to the assembly, with higher scores indicating a higher quality assembly. A score of 0.31 ranks this assembly higher than the majority of other published transcriptomes, with 87% of reads mapping, and only 6% of bases uncovered (no read support) and 8% contigs low-covered (mean per-base read coverage of < 10).

### Sequence Read Mapping and Estimation of Gene Expression

Raw sequencing reads corresponding to individual kidney samples were mapped to the reference kidney transcriptome using Salmon, which resulted in an average mapping rate of 85% (sd=4%). These mapping data were imported into R and summarized into gene-level counts using tximport, after which, edgeR was used to generate normalized estimates of gene expression.

### Global Analysis of Differential Gene Expression

Analysis of global patterns of gene expression, aimed at uncovering previously unknown differences in renal gene expression related to water deprivation by examining the entire transcriptome, was conducted using edgeR. After normalizing the count data using the TMM method (69), which is done by finding a set of scaling factors for the library sizes that minimize the log-fold changes between the samples for most genes, we controlled for over 13,600 multiple comparisons to yield a robust statistical analysis of differential gene expression. This analysis revealed a dramatic response to acute dehydration in gene expression, with 465 genes (Figure 1) differentially expressed (252 higher expression in acute dehydration, 213 lower expression in dehydration).

**Figure 1.**
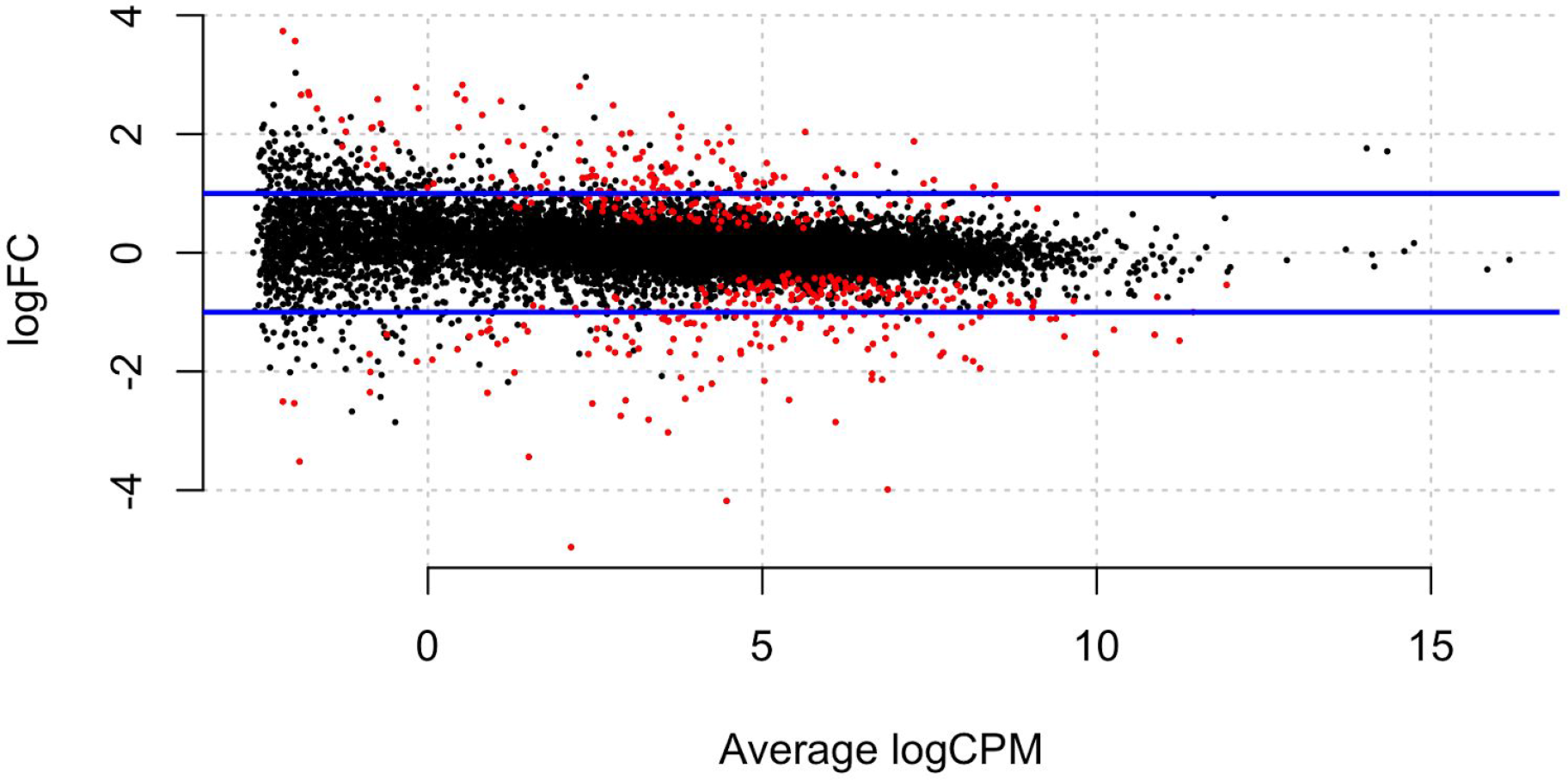
Smear plot of differential expression. Y-axis is the log fold change, where positive numbers indicate higher expression in control rodents, negative numbers indicate higher expression in acute dehydration. X-axis is the log counts per million, a metric of expression. Each dot is a gene, with red dots indicating significantly different gene expression (FDR < 0.01).

Gene ontology analysis of the 252 genes more highly expressed in acute dehydration (Table 3) suggests that acute dehydration is linked to first, the cellular starvation response, which occurs when cells are deprived of critical nutrients, including water. Significantly differentially expressed genes related to this term include ATF4 (Cyclic AMP-dependent transcription factor, known to interact with the endoplasmic reticulum (ER) stress pathway to promote the process of autophagy (70, 71)), SLC2A1 (Solute carrier family 2, facilitated glucose transporter member) and ATG14 (Beclin 1-associated autophagy-related key regulator), amongst others. Starvation stress does not seem to invoke widespread apoptosis (see below), which speaks to the ability of these mice to survive dehydration without resultant kidney damage.

Second, genes up-regulated in acute dehydration cluster around the GO term related to negative regulation of the MAPK cascade, which amongst other things, is an important regulator of transcription and translation (72) a phenomenon which is known to be a part of the response to hyperosmotic stress (27). Genes involved in this pathway, upregulated in the cactus mouse kidney include ERRFI1 (ERBB receptor feedback inhibitor 1), XBP1 (X-box-binding protein 1) and RGS2 (Regulator of G-protein signaling 2), the latter which has been shown to decrease the rate at which water is reabsorbed in the collecting tubules and duct via negative regulation of V2R signalling (73, 74).

Lastly, genes upregulated in acute dehydration are related to both positive and negative regulation of apoptosis. When cellular stress is prolonged or intense, and prosurvival mechanisms have failed, apoptotic pathways are invoked (75, 76). Genes involved in positive regulation include CTGF (Connective tissue growth factor), FOXO3 (Forkhead box protein O3), CASP8 (Caspase-8) and TRP53INP1 (Tumor protein p53-inducible nuclear protein 1), amongst others. In contrast, negative regulation of apoptosis could be important in mitigating the effects of cellular stress in an animal so highly adapted to dehydration. Genes related to this pathway include SOCS3 (Suppressor of cytokine signaling 3), CEBPB (CCAAT/enhancer-binding protein beta), PIM3 (Serine/threonine-protein kinase pim-3), EDN1 (Endothelin-1) and BCL2L1 (Bcl-2-like protein 1). Indeed, when looking globally, a strong signal of apoptosis is not observed. For instance, CHOP - the gene that connects the ER stress response to the apoptotic pathways (1, 70) is not differentially expressed (Figure 2), which suggests that the pro-survival mechanisms are largely successful. Additionally, key genes in the apoptotic pathway (*e.g.*, p53, BIM, BID, BAM, BAK, CASP2/3) are not differentially expressed, further suggesting that apoptosis is not widespread.

**Figure 2:**
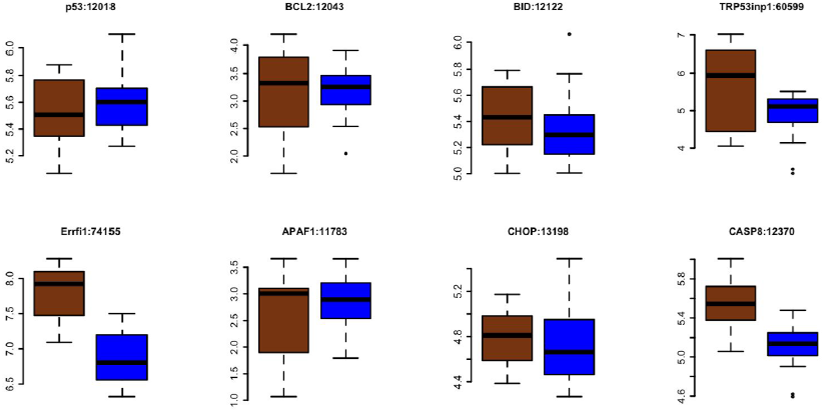
Apoptosis. In all cases, the Y-axis is the log transformed TPM. Brown boxplot refers to the acute dehydration group (n=17) while blue refers to the fully hydrated control animals (n=18). Botplot titles are of the format Gene Name:Entrez ID. The genes Trp53inp1, Errfi1, Casp8 are significantly differentially expressed.

**Table 3:** Top hierarchical level GO terms in the biological processes and cellular compartment categories for genes that are more highly expressed in acute dehydration.

| GO biological process | Term | Num genes | Num genes expected | Fold Enrichment | p-value |
| --- | --- | --- | --- | --- | --- |
|  | cellular response to starvation (GO:0009267) | 12 | 1.2 | 10.2 | 3.4E-05 |
|  | positive regulation of apoptotic process (GO:0043065) | 23 | 6.1 | 3.8 | 5.1E-04 |
|  | fatty acid metabolic process (GO:0006631) | 15 | 3.3 | 4.5 | 1.3E-02 |
|  | regulation of biological quality (GO:0065008) | 63 | 35.2 | 1.8 | 1.7E-02 |
|  | catabolic process (GO:0009056) | 34 | 14.2 | 2.4 | 1.8E-02 |
|  | negative regulation of apoptotic process (GO:0043066) | 25 | 8.9 | 2.8 | 3.3E-02 |
|  | small molecule biosynthetic process (GO:0044283) | 15 | 3.6 | 4.1 | 4.2E-02 |
|  | negative regulation of MAPK cascade (GO:0043409) | 10 | 1.6 | 6.3 | 4.8E-02 |
| **GO cellular component** | **Term** | **Num genes** | **Num genes expected** | **Fold Enrichment** | **p-value** |
|  | cytosol (GO:0005829) | 41 | 16.1 | 2.5 | 4.7E-05 |
|  | mitochondrion (GO:0005739) | 38 | 18.5 | 2.1 | 2.4E-02 |
|  | extracellular exosome (GO:0070062) | 50 | 27.4 | 1.8 | 2.4E-02 |
|  | organelle membrane (GO:0031090) | 36 | 16.7 | 2.2 | 1.6E-02 |

Gene ontology analysis of the 213 genes lowly expressed in acute dehydration (Table 4) suggests that acute dehydration is linked to lower expression in genes related to cholesterol biosynthetic processes. Genes related to this pathway include LSS (Lanosterol synthase), HMGCS1 (Hydroxymethylglutaryl-CoA synthase, cytoplasmic) and FDFT1 (Squalene synthase). Second, genes expressed significantly lower in the cactus mouse kidney during acute dehydration cluster around the GO term extracellular matrix organization. Genes involved in this pathway, and differentially expressed, include FN1 (Fibronectin), COL4A1, COL4A6, COL3A1, COL2A1, COL1A1 and COL1A2 (Collagen alpha chain genes) as well as LUM (Lumican). With regards to Fibronectin and Collagen, these proteins have been commonly associated with extracellular matrix turnover and renal interstitial fibrosis (7, 77). Renal interstitial fibrosis, a hallmark of kidney injury, has been causally linked to renal functional impairment (78), with fibrosis leading to a reduction in glomerular filtration rate (GFR) and overall function. That these genes are lowly expressed in the kidney of acutely dehydrated cactus mice provides strong evidence for the hypothesis that dehydration is not related to widespread kidney damage, and further, that extracellular matrix turnover is lowered.

**Table 4:** Top hierarchical level GO terms in the biological processes and cellular compartment categories for genes that are more lowly expressed in acute dehydration.

| **GO biological process** | **Term** | **Num genes** | **Num genes expected** | **Fold Enrichment** | **p-value** |
| --- | --- | --- | --- | --- | --- |
|  | cholesterol metabolic process (GO:0008203) | 15 | 0.9 | 17.4 | 1.9E-10 |
|  | isoprenoid biosynthetic process (GO:0008299) | 8 | 0.2 | 33.1 | 1.6E-06 |
|  | extracellular matrix organization (GO:0030198) | 14 | 1.6 | 8.8 | 1.0E-05 |
|  | positive regulation of bone mineralization (GO:0030501) | 6 | 0.3 | 18.8 | 8.4E-03 |
|  | cellular response to organic substance (GO:0071310) | 31 | 12.2 | 2.5 | 1.5E-02 |
|  | response to organonitrogen compound (GO:0010243) | 17 | 4.6 | 3.7 | 3.2E-02 |
|  | renal system development (GO:0072001) | 12 | 2.3 | 5.2 | 3.8E-02 |
|  | response to acid chemical (GO:0001101) | 11 | 2.0 | 5.6 | 5.0E-02 |
| **GO cellular component** | **Term** | **Num genes** | **Num genes expected** | **Fold Enrichment** | **p-value** |
|  | extracellular space (GO:0005615) | 41 | 13.1 | 3.1 | 8.4E-08 |
|  | fibrillar collagen trimer (GO:0005583) | 5 | 0.1 | 48.2 | 1.1E-04 |
|  | endoplasmic reticulum (GO:0005783) | 34 | 13.1 | 2.6 | 4.0E-04 |
|  | basement membrane<br>(GO:0005604) | 9 | 0.9 | 10.1 | 4.7E-04 |
|  | cell surface (GO:0009986) | 21 | 7.3 | 2.9 | 1.8E-02 |

### Critical Process Analysis

To better understand the response to acute dehydration in desert adapted animals, we analyzed patterns of differential expression between control animals and those exposed to acute dehydration for specific processes proposed to be critical to desert survival. To identify genes related to specific processes of interest (water balance and sodium regulation), we selected relevant gene ontology terms then identified genes in *Mus* that correspond to that term. Expression of the *Peromyscus* orthologs of these genes was estimated.

#### Water

For desert animals, the regulation of water is obviously critical (8, 14, 15). Here, the Aquaporin genes are thought to be critical to the maintenance of homeostasis, and as such have been the focus of study for decades (79, 80). How desert animals use these genes to regulate water is largely unstudied, though one study demonstrates that AQP4 is absent in at least one species of the *Dipodomys* rodents (19). The transcriptomic mechanisms *P. eremicus* uses to regulate water appear to depends critically on INPP5K (Figure 3), which is known to be expressed in the collecting duct, and is functionally responsible for urine concentration (81) and is involved in insulin signaling (82). Interestingly, this gene may also have a role in negatively regulating actin in the extracellular matrix (83), which provides yet another indication of the general depression of turnover in the extracellular matrix, as does the lower expression of Fibronectin and the Collagens.

In addition to the INPP5K gene, Aquaporin 4 (Figure 4) is significantly differentially expressed, being lower in acute dehydration relative to control animals. AQP4, expressed in the basolateral membrane of the collecting duct, is responsible for transport of water out of the cell, to the interstitium (84, 85). Defects in this non-vasopressin-responsive AQP may result in deficiencies in urine concentration (86). The functional correlates of lower expression are currently unknown. On one hand, lower expression might suggest a reduced capacity for moving water from membrane cells to the interstitium while on the other, in severe acute dehydration, very little water is available for movement, so perhaps there is little need to manufacture such protein complexes. Of note, no other Aquaporin was significantly differentially expressed in acute dehydration. This is perhaps surprisingly, especially for AQP2, whose role is predominately in reabsorption of water from the urinary lumen into the cell (84). Such RNAseq studies of acute dehydration have not been previously conducted, and as such, the complexity of the response has not been previously fully characterized. Ongoing work in the lab aims to provide further functional characterization of the responses to acute dehydration.

**Figure 3.**
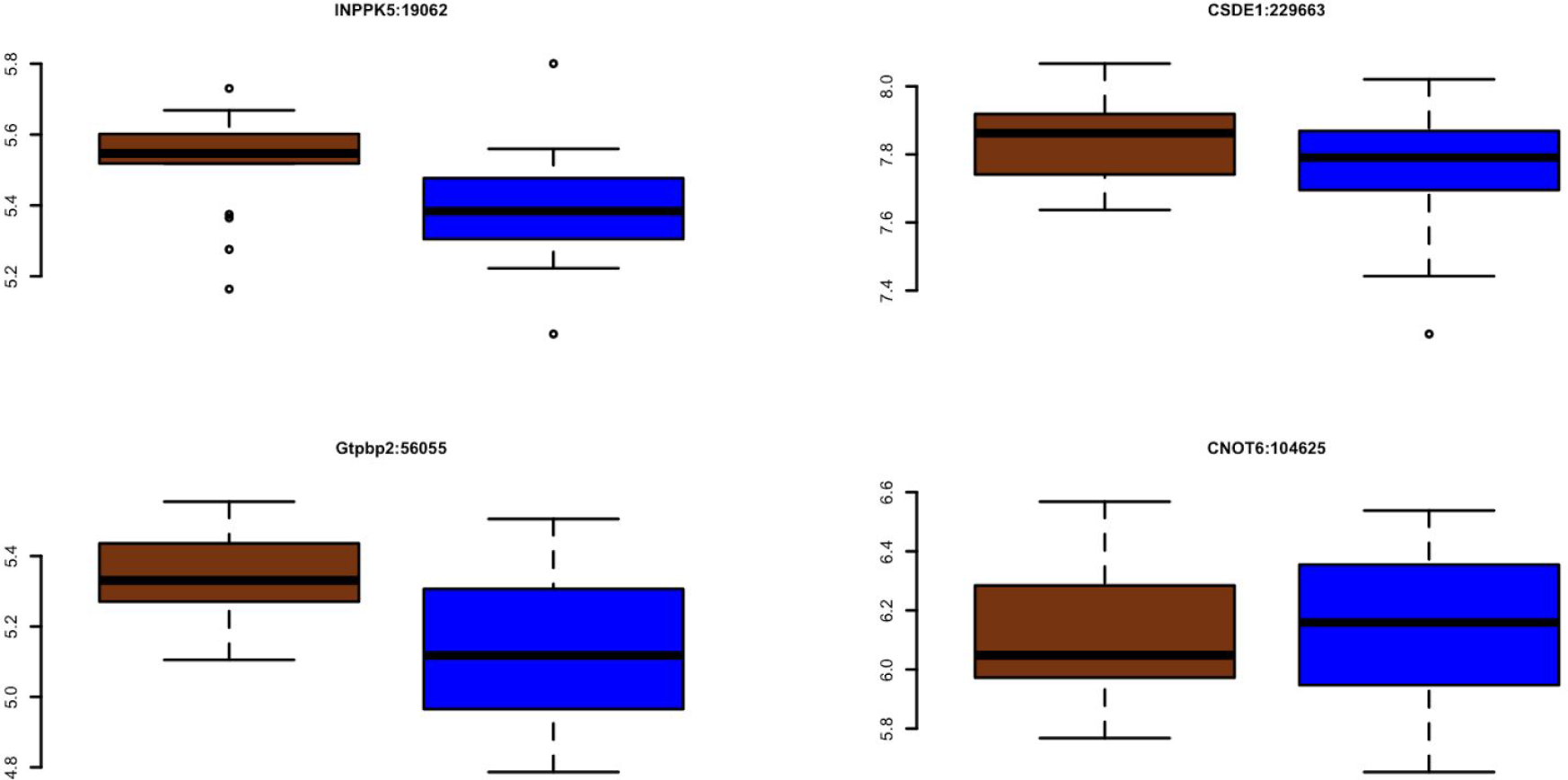
Regulation of renal water transport. In all cases, the Y-axis is the log transformed TPM. Brown boxplot refers to the acute dehydration group (n=17) while blue refers to the fully hydrated control animals (n=18). Botplot titles are of the format Gene Name:Entrez ID. INPP5K is significantly differentially expressed while the others are not.

**Figure 4.**
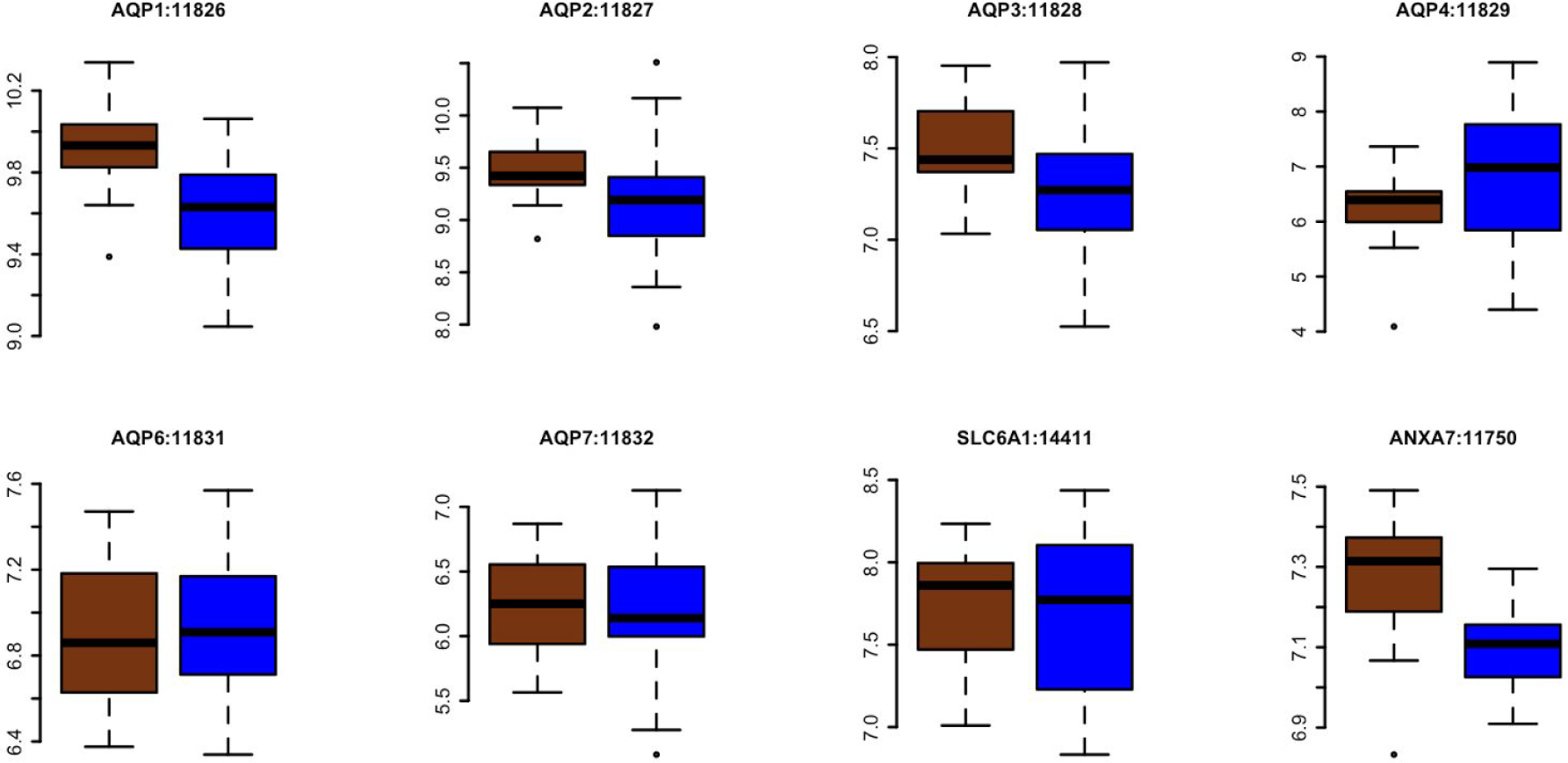
Cellular Water Homeostasis. In all cases, the Y-axis is the log transformed TPM. Brown boxplot refers to the acute dehydration group (n=17) while blue refers to the fully hydrated control animals (n=18). Botplot titles are of the format Gene Name:Entrez ID.

#### Sodium

In addition to the management of water, desert animals must also manage the maintenance of salts, namely Sodium. Indeed, hypernatremia is common in dehydration (26), and observed in acutely dehydrated cactus mice (64). In response to dehydration related hypernatremia, Avpr2, SLC8A1, Scnn1a, AGT and REN are significantly differentially expressed (Figure 5), with Avpr2, SLC8A1 being more highly expressed in control animals. Avpr2, a vasopressin receptor, has been linked to both sodium excretion (87) and the development of nephrogenic diabetes insipidus (88), while SLC8A1 is a sodium/calcium exchanger (89). Highly expressed in acute dehydration, Scnn1a is one component of the renal epithelial sodium channel (90) and had been linked to the development of pseudohypoaldosteronism (91, 92), a condition in which sodium is released inappropriately, resulting in hyponatremia. Lastly, angiotensinogen and renin, more highly expressed in acute dehydration, are thought to be broadly involved in the management of blood pressure and sodium excretion (93, 94). In summary, and in contrast to the relatively simple mechanisms through which water is regulated, the maintenance of serum Sodium levels are seemingly more complex, with multiple genes working in concert, in the face of hypernatremia, to regulate serum Sodium. Elucidating the individual roles or each gene uncovered here could be an exceptionally fruitful area of research. For instance, does each gene have a specific role in maintaining balance, or is there a high degree of redundancy in the regulation of serum Sodium.

**Figure 5:**
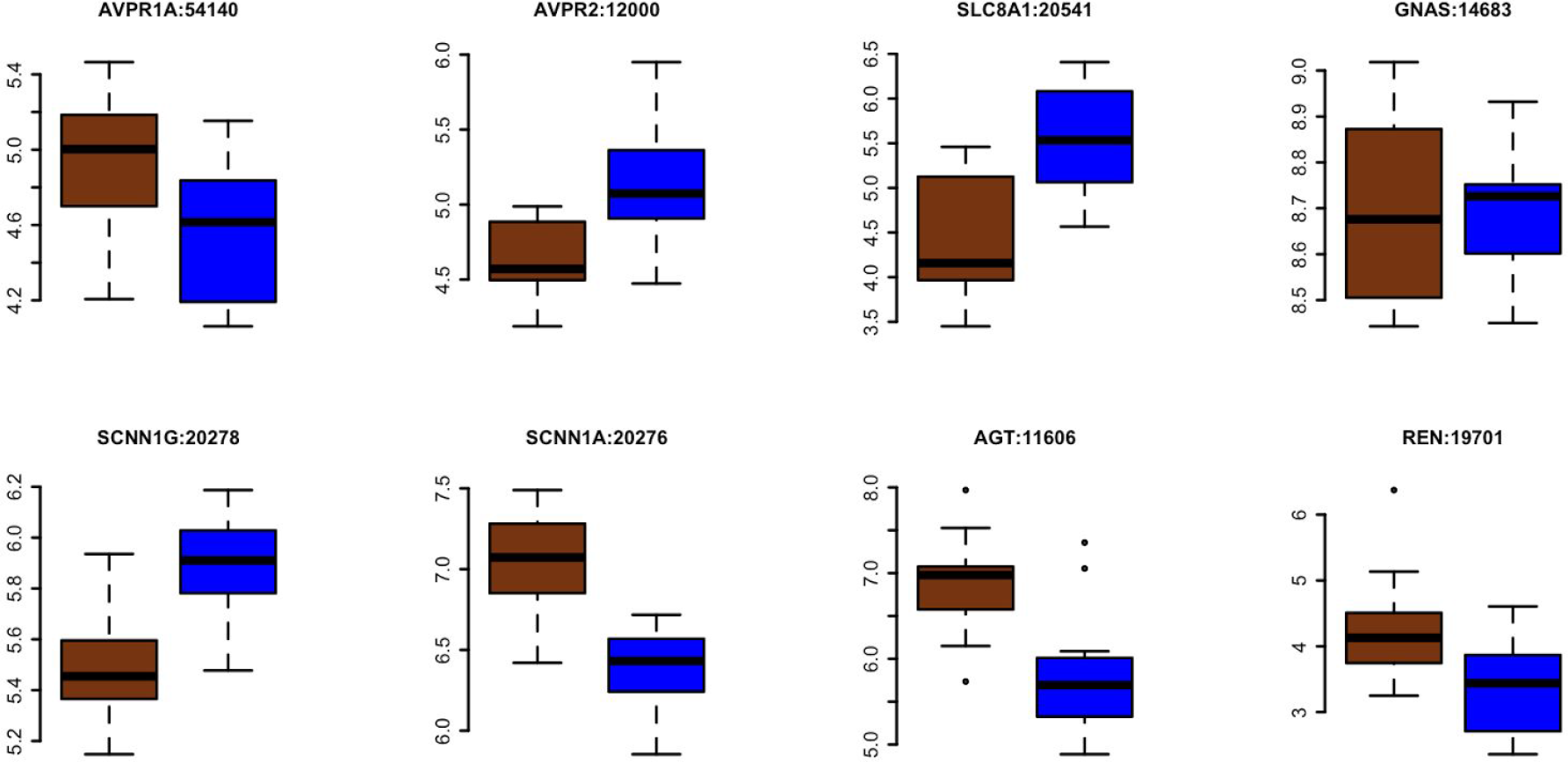
Sodium Ion Homeostasis. In all cases, the Y-axis is the log transformed TPM. Brown boxplot refers to the acute dehydration group (n=17) while blue refers to the fully hydrated control animals (n=18). Botplot titles are of the format Gene Name:Entrez ID. Avpr2, SLC8A1, Scnn1a, AGT and REN are significantly differentially expressed.

### Coexpression network analysis

WCGNA analysis identified 22 coexpression modules related to the measured phenotypes. While nearly all modules had significant relationships with at least one physiology measurement (Figure 6), several modules showed a particularly tight linkage with physiology. The gene ontology terms of each module are available in Supplementary table 2 (name = GOEnrichmentTable.csv on GitHub). The dehydration-linked weight loss of animals has a significant positive (higher gene expression related to more weight loss) relationship with the ‘tan’ module, which contains GO-terms transmembrane signaling receptor activity, circulatory and cardiovascular system development and peptide hormone binding. The three described genes mostly tightly linked to this module include CPM (carboxypeptidase M) which has, potentially, angiotensinogen converting function (95), SDK2 (sidekick cell adhesion molecule 2), and GALC (galactosylceramidase). Dehydration-linked weight loss of animals is has a significant negative relationship (higher gene expression related to less weight loss) with the ‘grey’ module, which contains GO-terms linked to monooxygenase activity, steroid metabolic process, and negative regulation of transport. The three described genes mostly tightly linked to this module include ITGA99 (integrin alpha 9), SGCG (sarcoglycan, gamma) and MAGBE4 (melanoma antigen, family B, 4). Broadly speaking, the integrins are receptors for extracellular matrix components (96) and are involved in the regulation in intracellular osmolarity (97). Expression of integrins, which could allow for sensing of osmotic stress and subsequent production of osmolytes, seem to have important links to dehydration-linked weight loss.

Sodium and Chloride ion concentrations were both positively associated (higher gene expression related to higher serum concentrations) with genes in the ‘royalblue’ module, which contained the GO terms vitamin transport, water transmembrane transporter activity, and water channel activity. Top genes in this group include CYP4B1 (cytochrome P450, family 4, subfamily b), GSTA1 (glutathione S-transferase, alpha 1), and LTA4H (leukotriene A4 hydrolase). Sodium and Chloride ion concentration has a negative relationship with genes in the grey module (see Dehydration-linked weight loss), suggesting the tight physiological link between water and electrolyte homeostasis.

**Figure 6:**
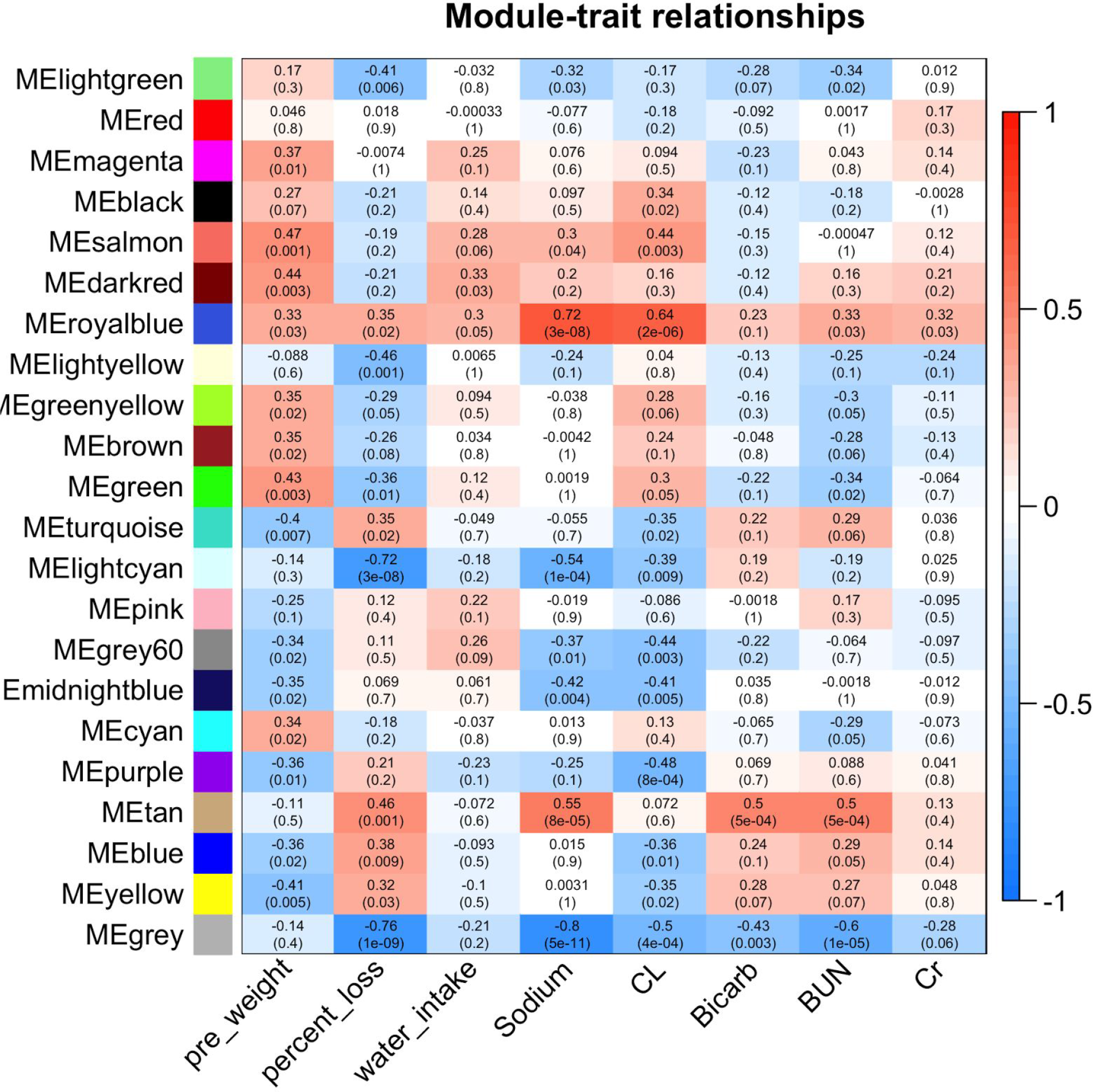
Heatmap describing the trait-module relationship that exist in the dataset. Colors on the lefthand axis represent the coexpression module, while the righthand axis represents the strength of the correlation. Within each cell, the top number represents the correlation coefficient while the lower number in parentheses is the p-value of the correlation.

### Biomarkers of kidney damage

A great deal of research has gone into identifying markers of acute kidney injury (1, 98–100), and as a result, the measurement of kidney damage has moved well beyond assaying serum Creatinine, which is known to be relatively insensitive and slow to indicate damage (101). In humans, dehydration and other clinical conditions that cause a pathological reduction in renal perfusion pressure are responsible for acute kidney injury (102). If acute dehydration results in substantial tissue damage in the cactus mouse, this damage will be evident when looking at a panel of biomarkers specific for kidney injury. Despite these markers not having been validated in the cactus mouse, that a large number of statistically independent tests are concordant (Figure 7) showing no differential expression, suggests that no significant tissue damage is occurring.

**Figure 7:**
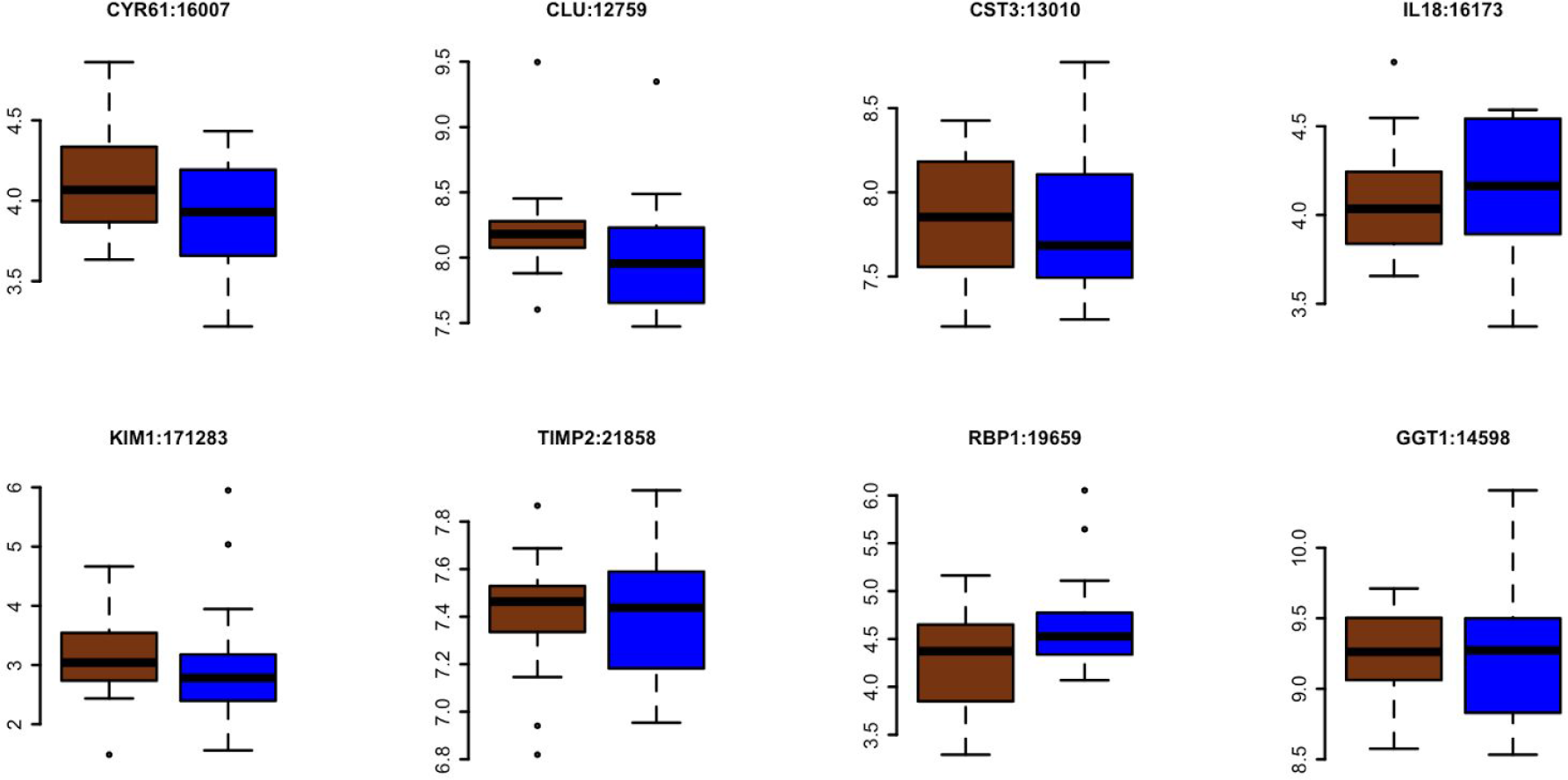
Biomarkers of acute kidney injury. In all cases, the Y-axis is the log transformed TPM. Brown boxplot refers to the acute dehydration group (n=17) while blue refers to the fully hydrated control animals (n=18). Botplot titles are of the format Gene Name:Entrez ID.

## Conclusion

Animal living in desert environments are forced to survive despite severe heat, intense solar radiation, and both acute and chronic dehydration. Indeed, these animals have evolved phenotypes that address these environmental stressors. To begin to understand the ways in which the desert adapted rodent *P. eremicus* survives, we performed an experiment by which we subjected reproductively mature adults to profound acute dehydration during which they lost, on average 23% of their body weight over the three day experiment. We modeled this loss after a scenario of a summertime rainfall event, followed by intense drought. While animals lost a substantial amount of weight, they were behaviorally intact. Animals react via a series of changes in the kidney, which include up-regulating genes responsible for 1. reducing the rate of transcription, and 2. Maintaining water and salt balance. Extracellular matrix turnover appears to be substantially decreased, and apoptosis appears to be limited. Serum Creatinine and other markers of kidney injury are not elevated over baseline, which is different than *Mus* exposed to less severe dehydration (7) and suggests that acute dehydration is not responsible for widespread kidney damage in the cactus mouse.

The work presented here are the results of experiment allowing us to peer deeply into the renal machinery of *P*. *eremicus* allowing for survival despite severe acute dehydration. In contrast, humans are exquisitely sensitive to alteration on water balance, with even minor dehydration resulting in physiological compromise (26). The molecular toolkit employed by the cactus mouse - the genes identified in this study - have orthologues in human. One exciting potential outcome of this line of work is the ability to develop manipulative genetic or pharmacologic techniques enabling a more robust human response to dehydration. Indeed, such a response could be extremely valuable in a wide variety of situation where potable water is limited.

## References

1 Zuk A, Bonventre JV (2016) Acute Kidney Injury. Annu Rev Med 67:293–307.

2 Mehta RL, et al. (2015) International Society of Nephrology’s 0by25 initiative for acute kidney injury (zero preventable deaths by 2025): a human rights case for nephrology. Lancet 385(9987):2616–2643.

3 IPCC (2007) Working Group I Report “The Physical Science Basis” (Cambridge Univ Press, Cambridge) Available at: http://www.ipcc.ch/publications_and_data/ar4/wg1/en/contents.html.

4 Roberts EM, Pope GR, Newson MJF, Lolait SJ, O’Carroll AM (2010) The Vasopressin V1b Receptor Modulates Plasma Corticosterone Responses to Dehydration-Induced Stress. J Neuroendocrinol 23(1):12–19.

5 Greenleaf JE(1992) http://www.rstudio.com/ Problem: thirst, drinking behavior, and involuntary dehydration. Med Sci Sports Exerc 24(6):645–656.

6 Bouby N, Fernandes S (2003) Mild dehydration, vasopressin and the kidney: animal and human studies. Eur J Clin Nutr 57 Suppl 2:S39–46.

7 Roncal Jimenez CA, et al. (2014) Fructokinase activity mediates dehydration-induced renal injury. Kidney Int 86(2):294–302.

8 Nagy KA, Gruchacz MJ (1994) Seasonal Water and Energy-Metabolism of the Desert-Dwelling Kangaroo Rat (Dipodomys merriami). Physiol Zool 67(6):1461–1478.

9 Takei Y, Bartolo RC, Fujihara H, Ueta Y, Donald JA (2012) Water deprivation induces appetite and alters metabolic strategy in Notomys alexis: unique mechanisms for water production in the desert. Proceedings Of The Royal Society B-Biological Sciences 279(1738):2599–2608.

10 MaresMA(1983) Desert rodent adaptation and community structure. Great Basin Naturalist Memoirs 0(7):30–43.

11 KaisslingB, De RouffignacC, Barrett JM, Kriz W (1975) The structural organization of the kidney of the desert rodent Psammomys obesus. Anat Embryol 148(2):121–143.

12 Gottschalk CW, et al. (1963) Micropuncture Study of Composition of Loop of Henle Fluid in Desert Rodents. Am J Physiol 204(4):532–535.

13 Altschuler EM, Nagle RB, Braun EJ, Lindstedt SL, Krutzsch PH (1979) Morphological study of the desert heteromyid kidney with emphasis on the genus Perognathus. Anat Rec 194(3):461–468.

14 Schmidt-Nielsen K, Schmidt-Nielsen B (1952) Water metabolism of desert mammals 1. Physiol Rev 32(2):135–166.

15 Cortés A, Rosenmann M, Bozinovic F (2000) Water economy in rodents: evaporative water loss and metabolic water production. Rev Chil Hist Nat 73(2):311–321.

16 Ma TH, et al. (1998) Severely impaired urinary concentrating ability in transgenic mice lacking aquaporin-1 water channels. J Biol Chem 273(8):4296–4299.

17 Brown D, Katsura T, Kawashima M, Verkman AS, Sabolic I (1995) Cellular distribution of the aquaporins: A family of water channel proteins. Histochem Cell Biol 104(1):1–9.

18 Verkman AS (2002) Physiological importance of aquaporin water channels. Ann Med 34(3):192–200.

19 Huang Y, et al. (2001) Absence of Aquaporin-4 water channels from kidneys of the desert Rodent Dipodomys merriami merriami.American Journal Of Physiology-Renal Physiology 280(5):F794–F802.

20 Marra NJ, Romero A, DeWoody A (2014) Natural selection and the genetic basis of osmoregulation in heteromyid rodents as revealed by RNA-seq. Mol Ecol 23(11):2699–2711.

21 Marra NJ, Eo SH, Hale MC, Waser PM, DeWoody A (2012) A priori and a posteriori approaches for finding genes of evolutionary interest in non-model species: Osmoregulatory genes in the kidney transcriptome of the desert rodent Dipodomys spectabilis (Banner-Tailed Kangaroo Rat). Comparative Biochemistry and Physiology - Part D: Genomics and Proteomics:1–12.

22 Giorello FM, et al. (2014) Characterization of the kidney transcriptome of the South American olive mouse Abrothrix olivacea. BMC Genomics 15(1):446.

23 Esteva-Font C, et al. (2012) Molecular biology of water and salt regulation in the kidney. Cell Mol Life Sci 69(5):683–695.

24 MacManes MD, Eisen MB (2014) Characterization of the transcriptome, nucleotide sequence polymorphism, and natural selection in the desert adapted mouse Peromyscus eremicus. PeerJ 2:e642.

25 Hediger MA, et al. (2004) The ABCs of solute carriers: physiological, pathological and therapeutic implications of human membrane transport proteins. Pflugers Arch 447(5):465–468.

26 Thomas DR, et al. (2008) Understanding clinical dehydration and its treatment. J Am Med Dir Assoc 9(5):292–301.

27 Burg MB, Ferraris JD, Dmitrieva NI (2007) Cellular response to hyperosmotic stresses. Physiol Rev 87(4):1441–1474.

28 Ferraris JD, Williams CK, Ohtaka A, García-Pérez A (1999) Functional consensus for mammalian osmotic response elements. Am J Physiol 276(3 Pt 1):C667–73.

29 Kumar A, Tyagi MG (2011) Hypertonicity induced modulation of gene transcription and translation of water regulatory molecules. Biomed Res 22(1):93–101.

30 Michea L, et al. (2000) Cell cycle delay and apoptosis are induced by high salt and urea in renal medullary cells. Am J Physiol Renal Physiol 278 (2):F209–18.

31 Kültz D, Chakravarty D (2001) Hyperosmolality in the form of elevated NaCl but not urea causes DNA damage in murine kidney cells. Proc Natl Acad Sci U S A 98(4):1999–2004.

32 Dmitrieva NI, Bulavin DV, Burg MB (2003) High NaCl causes Mre11 to leave the nucleus, disrupting DNA damage signaling and repair. Am J Physiol Renal Physiol 285(2):F266–74.

33 Zhou X, Ferraris JD, Cai Q, Agarwal A, Burg MB (2005) Increased reactive oxygen species contribute to high NaCl-induced activation of the osmoregulatory transcription factor TonEBP/OREBP. Am J Physiol Renal Physiol 289(2):F377–85.

34 Robbins E, Pederson T, Klein P (1970) Comparison of mitotic phenomena and effects induced by hypertonic solutions in HeLa cells. J Cell Biol 44(2):400–416.

35 Santos BC, Chevaile A, Kojima R, Gullans SR (1998) Characterization of the Hsp110/SSE gene family response to hyperosmolality and other stresses. Am J Physiol 274(6 Pt 2):F1054–61.

36 Hampton RY (2000) ER stress response: getting the UPR hand on misfolded proteins. Curr Biol 10(14):R518–21.

37 Veal R, Caire W (1979) Peromyscus eremicus. Mammalian Species:1–6.

38 Drickamer LC, Vestal BM (1973) Patterns of reproduction in a laboratory colony of Peromyscus. J Mammal 54(2):523–528.

39 Dewey MJ, Dawson WD (2001) Deer mice: “The Drosophila of North American mammalogy.” Genesis 29(3):105–109.

40 Bester-Meredith JK, Marler CA (2007) Social Experience During Development and Female Offspring Aggression in Peromyscus Mice. Ethology 113(9):889–900.

41 Dunmire WW (1960) An Altitudinal Survey of Reproduction in Peromyscus maniculatus. Ecology 41(1):174–182.

42 Foltz DW (1981) Genetic evidence for long-term monogamy in a small rodent, Peromyscus polionotus. Am Nat 117(5):665–675.

43 Steiner CC, Weber JN, Hoekstra HE (2007) Adaptive variation in beach mice produced by two interacting pigmentation genes. PLoS Biol 5(9):e219.

44 Mullen LM, Hoekstra HE (2008) Natural selection along an environmental gradient:a classic cline in mouse pigmentation. Evolution 62(7):1555–1570.

45 Linnen CR, et al. (2009) On the origin and spread of an adaptive allele in deer mice. Science 325(5944):1095–1098.

46 Kordonowy LL, MacManes MD (2016) Characterization of a male reproductive transcriptome for Peromyscus eremicus (Cactus mouse). PeerJ 4:e2617.

47 Sikes RS, Gannon WL, Animal Care and Use Committee of the American Society of Mammalogists (2011) Guidelines of the American Society of Mammalogists for the use of wild mammals in research. J Mammal 92(1):235–253.

48 Andrews SR (2016) FastQC Available at: http://www.bioinformatics.babraham.ac.uk/projects/fastqc/.

49 MacManes MD (2015) Establishing evidenced-based best practice for the de novo assembly and evaluation of transcriptomes from non-model organisms. biorxiv.org (bioinformatics):1–23.

50 Song L, Florea L (2015) Rcorrector: efficient and accurate error correction for Illumina RNA-seq reads. Gigascience 4(1):48.

51 Jiang H, Lei R, Ding S-W, Zhu S (2014) Skewer: a fast and accurate adapter trimmer for next-generation sequencing paired-end reads. BMC Bioinformatics 15(1):182.

52 Haas BJ, et al. (2013) De novo transcript sequence reconstruction from RNA-seq using the Trinity platform for reference generation and analysis. Nat Protoc 8(8):1494–1512.

53 Liu J, et al. (2016) BinPacker: Packing-Based De Novo Transcriptome Assembly from RNA-seq Data. PLoS Comput Biol 12(2):e1004772.

54 Kannan S, Hui J, Mazooji K, Pachter L, Tse D (2016) Shannon: An Information-Optimal de Novo RNA-Seq Assembler (Cold Spring Harbor Labs Journals) Available at: http://biorxiv.org/lookup/doi/10.1101/039230.

55 Simão FA, Waterhouse RM, Ioannidis P, Kriventseva EV, Zdobnov EM (2015) BUSCO: assessing genome assembly and annotation completeness with single-copy orthologs. Bioinformatics 31(19):3210–3212.

56 Smith-Unna RD, Boursnell C, Patro R, Hibberd JM, Kelly S (2015) TransRate: reference free quality assessment of de-novo transcriptome assemblies. bioRxiv:1–25.

57 Patro R, Duggal G, Kingsford C (2015) Accurate, fast, and model-aware transcript expression quantification with Salmon. biorxiv.org:1–35.

58 Camacho C, et al. (2009) BLAST+: architecture and applications. BMC Bioinformatics 10(1):421.

59 RStudio Team (2015) RStudio: Integrated Development for R. Available at: http://www.rstudio.com/ [Accessed September 1, 2016].

60 Soneson C, Love MI, Robinson MD (2015) Differential analyses for RNA-seq: transcript-level estimates improve gene-level inferences. F1000Res 4:1521–1519.

61 Robinson MD, McCarthy DJ, Smyth GK (2010) edgeR: a Bioconductor package for differential expression analysis of digital gene expression data. Bioinformatics 26(1):139–140.

62 Young IT (1977) Proof without prejudice: use of the Kolmogorov-Smirnov test for the analysis of histograms from flow systems and other sources. J Histochem Cytochem 25(7):935–941.

63 Alexa A, Rahnenfuhrer J (2010) topGO: enrichment analysis for gene ontology. R package version 2(0). Available at: http://bioconductor.uib.no/2.7/bioc/html/topGO.html.

64 Kordonowy L, et al. (2016) Physiological and biochemical changes associated with experimental dehydration in the desert adapted cactus mouse, Peromyscus eremicus. bioRxiv:047704.

65 Langfelder P, Horvath S (2008) WGCNA: an R package for weighted correlation network analysis. BMC Bioinformatics 9(1):559.

66 Waterhouse RM, Tegenfeldt F, Li J, Zdobnov EM, Kriventseva EV (2013) OrthoDB: a hierarchical catalog of animal, fungal and bacterial orthologs. Nucleic Acids Res 41(Database issue):D358–65.

67 Saier MHJr, Tran CV, Barabote RD (2006) TCDB: the Transporter Classification Database for membrane transport protein analyses and information. Nucleic Acids Res 34(Database issue):D181–6.

68 Griffiths-Jones S, et al. (2005) Rfam: annotating non-coding RNAs in complete genomes. Nucleic Acids Res 33(Database issue):D121–4.

69 McCarthy DJ, Chen Y, Smyth GK (2012) Differential expression analysis of multifactor RNA-Seq experiments with respect to biological variation. Nucleic Acids Res 40(10):4288–4297.

70 B’chir W, et al. (2013) The eIF2α/ATF4 pathway is essential for stress-induced autophagy gene expression. Nucleic Acids Res 41(16):7683–7699.

71 Darling NJ, Cook SJ (2014) The role of MAPK signalling pathways in the response to endoplasmic reticulum stress. Biochim Biophys Acta 1843(10):2150–2163.

72 Pende M, et al. (2004) S6K1–/–/S6K2–/– Mice Exhibit Perinatal Lethality and Rapamycin-Sensitive 5′-Terminal Oligopyrimidine mRNA Translation and Reveal a Mitogen-Activated Protein Kinase-Dependent S6 Kinase Pathway. Mol Cell Biol 24(8):3112–3124.

73 Zuber AM, Singer D, Penninger JM, Rossier BC, Firsov D (2007) Increased renal responsiveness to vasopressin and enhanced V2 receptor signaling in RGS2-/- mice. J Am Soc Nephrol 18(6):1672–1678.

74 Park F (2015) Accessory proteins for heterotrimeric G-proteins in the kidney. Front Physiol 6:219.

75 Kerr JF, Wyllie AH, Currie AR (1972) Apoptosis: a basic biological phenomenon with wide-ranging implications in tissue kinetics. Br J Cancer 26(4):239–257.

76 Fulda S, Gorman AM, Hori O, Samali A (2010) Cellular stress responses: cell survival and cell death. Int J Cell Biol 2010:214074.

77 Eddy AA (1996) Molecular insights into renal interstitial fibrosis. J Am Soc Nephrol 7(12):2495–2508.

78 Schainuck LI, Striker GE, Cutler RE, Benditt EP (1970) Structural-functional correlations in renal disease: Part II: The Correlations - ScienceDirect. Available at:http://www.sciencedirect.com/science/article/pii/S0046817770800612 [Accessed January 23, 2017].

79 van Os CH (1998) Role of aquaporins in renal water handling: physiology and pathophysiology. Nephrol Dial Transplant. Available at: http://ndt.oxfordjournals.org/content/13/7/1645.short.

80 Kwon T-H, et al. (2009) Aquaporins in the kidney. Handb Exp Pharmacol 190(190):95–132.

81 Pernot E, et al. (2011) The inositol Inpp5k 5-phosphatase affects osmoregulation through the vasopressin-aquaporin 2 pathway in the collecting system. Pflugers Arch 462(6):871–883.

82 Bridges D, Saltiel AR (2015) Phosphoinositides: Key modulators of energy metabolism. Biochim Biophys Acta 1851(6):857–866.

83 Balla T (2013) Phosphoinositides: tiny lipids with giant impact on cell regulation. Physiol Rev 93(3):1019–1137.

84 Nielsen S, et al. (2002) Aquaporins in the kidney: from molecules to medicine. Physiol Rev 82(1):205–244.

85 Holmes RP (2012) The role of renal water channels in health and disease. Mol Aspects Med. Available at: http://eutils.ncbi.nlm.nih.gov/entrez/eutils/elink.fcgi?dbfrom=pubmed&id=22252122&retmode=ref&cmd=prlinks.

86 Ma T, et al. (1997) Generation and phenotype of a transgenic knockout mouse lacking the mercurial-insensitive water channel aquaporin-4. J Clin Invest 100(5):957–962.

87 Bankir L, Fernandes S, Bardoux P, Bouby N, Bichet DG (2005) Vasopressin-V2 receptor stimulation reduces sodium excretion in healthy humans. J Am Soc Nephrol 16(7):1920–1928.

88 Neocleous V, et al. (2012/7) Identification and characterization of a novel X-linked AVPR2 mutation causing partial nephrogenic diabetes insipidus: A case report and review of the literature. Metabolism 61(7):922–930.

89 Gabellini N, Bortoluzzi S, Danieli GA, Carafoli E (2002) The human SLC8A3 gene and the tissue-specific Na+/Ca2+ exchanger 3 isoforms. Gene 298(1):1–7.

90 Christensen BM, et al. (2010) Sodium and potassium balance depends on αENaC expression in connecting tubule. J Am Soc Nephrol 21(11):1942–1951.

91 Dirlewanger M, et al. (2011) A homozygous missense mutation in SCNN1A is responsible for a transient neonatal form of pseudohypoaldosteronism type 1. Am J Physiol Endocrinol Metab 301(3):E467–73.

92 Welzel M, et al. (2013) Five novel mutations in the SCNN1A gene causing autosomal recessive pseudohypoaldosteronism type 1. Eur J Endocrinol 168(5):707–715.

93 Gociman B, et al. (2004) Expression of angiotensinogen in proximal tubule as a function of glomerular filtration rate. Kidney Int 65(6):2153–2160.

94 Ingelfinger JR, Pratt RE, Ellison K, Dzau VJ (1986) Sodium regulation of angiotensinogen mRNA expression in rat kidney cortex and medulla. J Clin Invest 78(5):1311–1315.

95 Donoghue M, et al. (2000) A Novel Angiotensin-Converting Enzyme–Related Carboxypeptidase (ACE2) Converts Angiotensin I to Angiotensin 1-9. Circ Res 87(5):e1–e9.

96 Pozzi A, Zent R (2003) Integrins: sensors of extracellular matrix and modulators of cell function. Nephron Exp Nephrol 94(3):e77–84.

97 Moeckel GW, et al. (2006) Role of integrin α1β1 in the regulation of renal medullary osmolyte concentration. American Journal of Physiology - Renal Physiology 290(1):F223–F231.

98 Obeidat MA, et al. (2011) Post-transplant nuclear renal scans correlate with renal injury biomarkers and early allograft outcomes. Nephrol Dial Transplant 26(9):3038–3045.

99 Schrezenmeier EV, Barasch J, Budde K, Westhoff T, Schmidt-Ott KM (2016) Biomarkers in acute kidney injury–pathophysiological basis and clinical performance. Acta Physiol. Available at: http://onlinelibrary.wiley.com/doi/10.1111/apha.12764/full.

100 Goodsaid FM, et al. (2009) Novel Biomarkers of Acute Kidney Toxicity. Clinical Pharmacology & Therapeutics 86(5):490–496.

101 Baum N, Dichoso CC, Carlton CE (1975) Blood urea nitrogen and serum creatinine. Physiology and interpretations. Urology 5(5):583–588.

102 Van Biesen W, et al. (2005) Relationship between fluid status and its management on acute renal failure (ARF) in intensive care unit (ICU) patients with sepsis: a prospective analysis. J Nephrol 18(1):54–60.

